# Exogenous expression of a histone H3.3 isoform causes extranuclear divisions in the *C. elegans* intestine

**DOI:** 10.1101/2025.04.10.648275

**Authors:** Carmen Herrera Sandoval, Christopher Borchers, Kara L. Osburn, Becky Boyd, Scott T. Aoki

**Author notes:** These authors contributed equally (Carmen Herrera Sandoval, Christopher Borchers). Corresponding author: S.T. Aoki.

## Abstract

Histone proteins condense DNA into chromatin and play significant roles in gene regulation. Mutations that alter histone function can disrupt critical gene expression programs and are implicated in driving cancer and other human diseases. Most mechanistic research on histones has been conducted in cell culture, mitotic tissues, or animal disease models rather than in normal tissue. *Caenorhabditis elegans* is a model organism with a well-established developmental program and conserved histone mechanisms shared with other metazoans. Prior work generated several single-copy histone reporters expressing exogenous histone protein in the intestine. Surprisingly, the histone H3.3 reporter worms possessed extra nuclei in the anterior intestine. This phenotype was incompletely penetrant and arose early during larval development. Mutations predicted to perturb histone H3.3 reporter expression or deposition into chromatin ameliorated the phenotype. DNA Fluorescence In situ Hybridization (FISH) was used to measure chromosome copy number and revealed approximately half the DNA content in the extra nuclei versus undivided controls, suggesting aberrant nuclear division during normal rounds of intestinal endoreduplication. Thus, these reporters serendipitously identified a distinct function of histone H3.3 in triggering aberrant nuclear division during animal development. Other histone isoforms may have previously unrecognized biological functions in terminally differentiated tissue alongside their roles in mitotic tissue.

## Introduction

Nucleosomes are histone protein octamers that condense DNA into chromatin (1-3). The octamers assemble from H2A, H2B, H3, and H4 histone proteins (2-4). Histone subtypes and isoforms possess sequence variance that facilitate specialized functions in gene regulation (5). For example, histone H3.1 of the H3 family is recognized by the CAF-1 histone chaperone (6), has replication-dependent deposition (7, 8), and is typically associated with gene repression (9, 10). In contrast, histone H3.3 is recognized by HIRA (7) and DAXX-ATRX (11) chaperone complexes, has replication-independent deposition (7, 12), and is associated with gene expression (7, 12). Mutations in H3.3 is implicated in rare human cancers such as glioblastoma, osteosarcoma, and chondroblastoma (13). Prominent mutations such as H3.3(K27M) or H3.3(K36M) alter methylation and acetylation states of chromatin and consequently affect differentiation and cell fate decisions by disrupting canonical gene regulatory pathways (13). Thus, mutations that affect H3.3 function can have profound impacts on animal biology.

Prior work created histone reporters to visualize nuclei of the *C. elegans* intestine (14). At hatching, the intestine is fully formed with 20 cells and 20 nuclei organized in a linear, pair-wise fashion (14, 15). Late in the first larval stage, specific nuclei divide, resulting in 30-34 nuclei that serve the animal throughout its entire lifetime (14, 15). The intestine expands in size throughout larval development with intestinal nuclei undergoing endoreduplication, or repeated rounds of DNA replication without mitosis or cytokinesis, leading to large, polyploid nuclei (16). The intestinal reporters expressed histones representative of H2B, H3.1, and H3.3 isoforms (17). Surprisingly, histone H3.3 reporter animals possessed extra anterior intestinal nuclei. Histones H3.1 and H3.3 vary by 4 amino acids in humans (18) and 9 amino acids in the *C. elegans* nematode (19). This high level of conservation stimulated investigation of how these modest histone isoform differences caused an unexpected phenotype in an animal model.

## Results and Discussion

### *his-72*/H3.3 reporter worms have extra anterior intestinal nuclei that arises early in larval development

*C. elegans* adult worms possess a set number of nuclei at the anterior region of the intestine **(Fig 1A)** (14). Prior work created intestinal HaloTag histone reporters (17) that consisted of an intestinal *elt-2* promoter (20, 21) driving histones H3.1 (*his-9)* or H3.3 (*his-72)* linked to a HaloTag (22) **(Fig 1B)**. As expected, both the histone H3.1 and H3.3 variants were expressed specifically in intestinal nuclei **(Fig 1C)**. Unexpectedly, imaging of reporter worms revealed *his-72*/H3.3 worms possess extra nuclei in the anterior portion of the intestine **(Fig 1C, D)**. *his-72*/H3.3 adult worms often possessed 7 or 8 nuclei in the first 6 cells, while *his-9*/H3.1 adult worms had primarily 6 nuclei **(Fig 1C, D)**. Analysis of nuclei number across the intestine revealed extranuclear divisions were restricted to the anterior region **(Fig S1A)**. Thus, some aspect of the *his-72*/H3.3 reporter worms caused extra nuclei to form sporadically in a specific region of the intestine.

**Fig 1.**
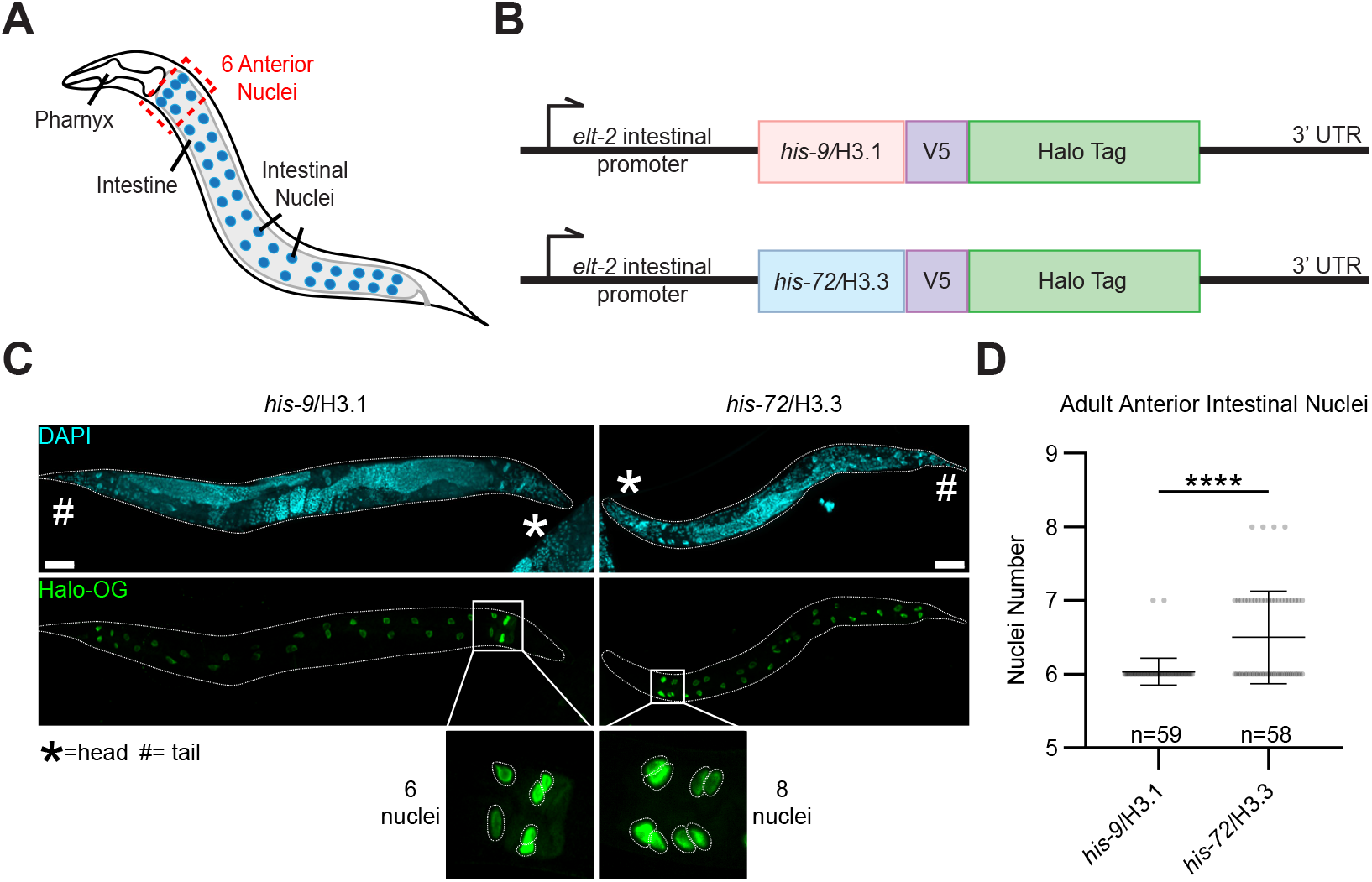
Histone H3.3 reporter worms exhibit an extra intestinal nuclei phenotype. (**A**) Nuclei organization in the intestine of *C. elegans*. An adult intestine normally possesses 20 cells with 30-34 nuclei (34 shown) with 6 anterior nuclei (dashed box). (**B**) Linear diagram of the HaloTag intestinal histone reporter. An *elt-2* intestine-specific promoter drives histone variants fused to a V5 epitope and HaloTag in *C. elegans*. Reporters also have an *unc-54* 3’UTR and intergenic region. (**C**) Fluorescent images of *his-9*/H3.1 and *his-72*/H3.3 reporter adult worms. Images shown as a combined montage max intensity projection of confocal stack taken at 40X magnification under oil immersion. Anterior nuclei shown at 3x magnification from original image. Scale bar, 50 µm. (**D**) Quantification of nuclei number in first 6 cells of anterior portion of intestine in adult reporter worms. Significance determined by unpaired student t-test; ****p<0.0001.

The *C. elegans* intestine has a well-established developmental pattern. During the first larval stage, the first six nuclei neither replicate their DNA or divide, the middle ten nuclei always replicate and divide, and the last four nuclei may or may not replicate and divide (14). Thus, a newly hatched L1 larva has 20 cells with 20 nuclei while a late L1 has 20 cells with 30 to 34 nuclei (14, 16). The number of intestinal cells and nuclei are maintained but grow as the worm develops. To determine when the extra nuclei phenotype arose, *his-72*/H3.3 and *his-9*/H3.1 reporter worms were collected as early L1 larvae or allowed to develop for 24 or 48 hours prior to fixing, staining, and imaging. Early L1 larvae of both *his-72*/H3.3 and *his-9*/H3.1 reporter worms had 6 anterior intestinal nuclei **(Fig 2A, C, D)**. By 24 hours into development, *his-72*/H3.3 reporter worms had more nuclei than the control *his-9*/H3.1 worms **(Fig 2B-D; Fig S1B)**. Aging in *C. elegans* plays a role in intestine function and is reported to affect nuclei number and integrity (23). To test whether aging exacerbated the extra nuclei phenotype, reporter worms were grown for 72 hours past adulthood, 6 days past L1, before collection and imaging. While analysis of nuclear counts at different ages revealed some differences in *his-72*/H3.3 worms (L1+24 hours and adults), no significant differences were observed across the remaining time points **(Fig 2D; Fig S1B, C)**. Extended aging did not induce additional intestinal nuclear divisions. These findings suggest that intestinal extranuclear division occurs post-embryonically at a very early stage in larval development.

**Fig 2.**
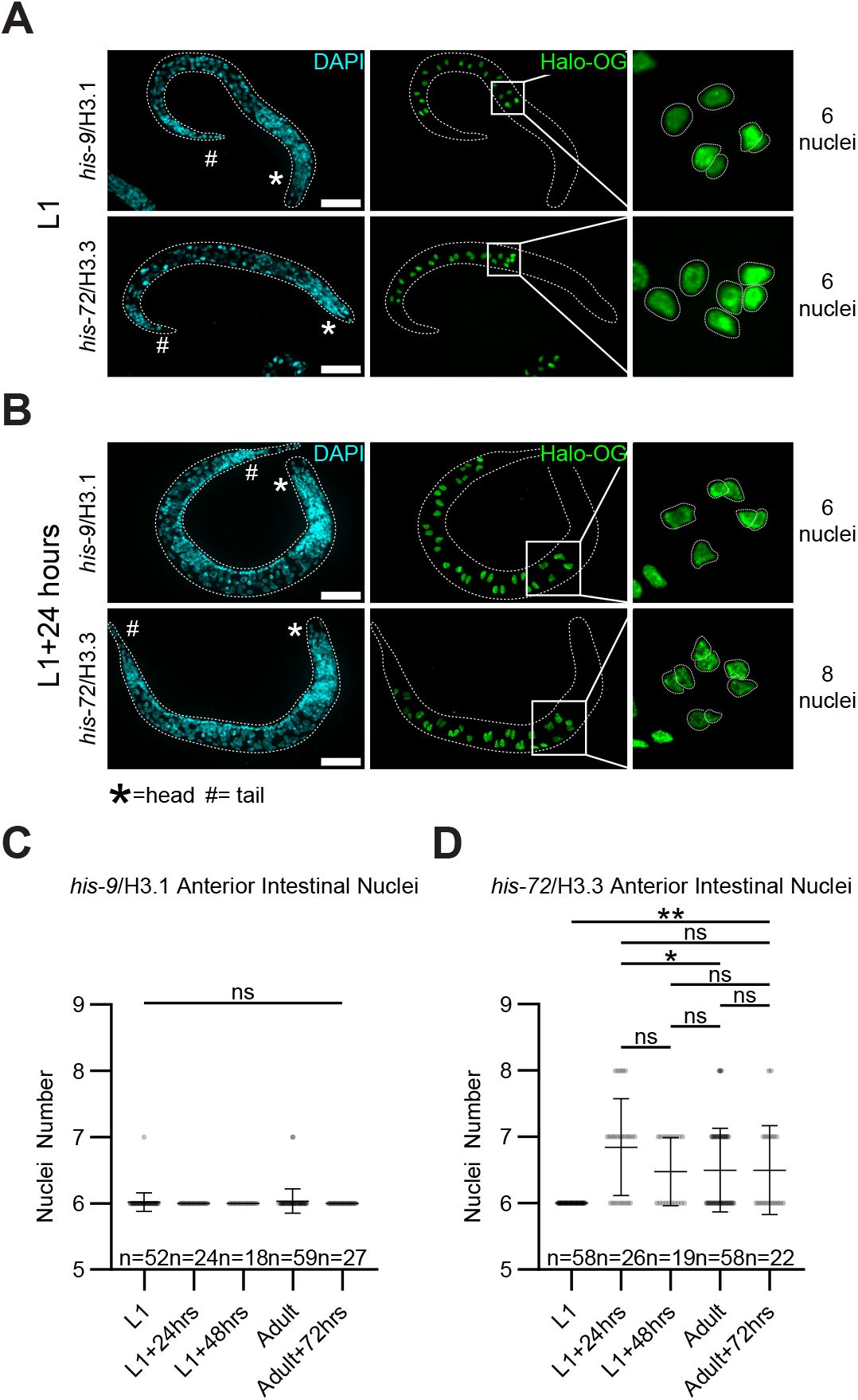
Intestinal extra nuclei phenotype arises during early development. (**A-B**) Fluorescent images of *his-9*/H3.1 and *his-72*/H3.3 reporter worms at the first larval stage (A) or grown for 24 hours (B). Images shown as max intensity projection of confocal stack taken at 63X magnification under oil immersion. Anterior nuclei shown at either 5x (L1) or 3x (L1+24 hours) magnification from original image. Scale bar, 10 µm. (**C-D**) Quantification of nuclei in anterior six intestinal cells in *his-9*/H3.1 (C) and *his-72*/H3.3 (D) reporter worms at various developmental stages. Adult data shown in Fig 1D used in (C) and (D). Significance was determined by performing a one-way ANOVA with Tukey’s multiple comparisons test; *p<0.05; **p<0.01; ns, not significant.

Temperature-induced environmental stress can impact *C. elegans* development (24-26). Histone reporter worms were propagated at different temperatures to evaluate their response to temperature stress. Lower temperature did not impact the extranuclear phenotype in *his-72*/H3.3 worms **(Fig S1D)**. Higher temperature could not be properly assessed due to observed worm lethality. In sum, the results indicated that the extra nuclei phenotype occurred during L1 larval development, but the divisions did not continue through later larval stages into adulthood, regardless of temperature conditions.

### *his-72*/H3.3 mutations ameliorate the intestinal extra nuclei phenotype

One concern was whether the extranuclear divisions observed were dependent on the *his-72* reporter. To test the necessity of reporter protein expression for the phenotype, CRISPR/Cas9 was used to create a frameshift mutation that introduced a premature stop codon in the reporter sequence **(Fig 3A**, see **Methods)**. The *his-72*/H3.3 frameshift reporter worms lacked expression of the reporter, as measured by imaging and immunoblot **(Figure S1E, F)**, and exhibited wild type nuclei numbers in adults **(Fig 3C)**. The results are supportive of the *his-72*/H3.3 reporter causing the observed intestinal nuclei phenotype rather than a random acquired mutation.

**Fig 3.**
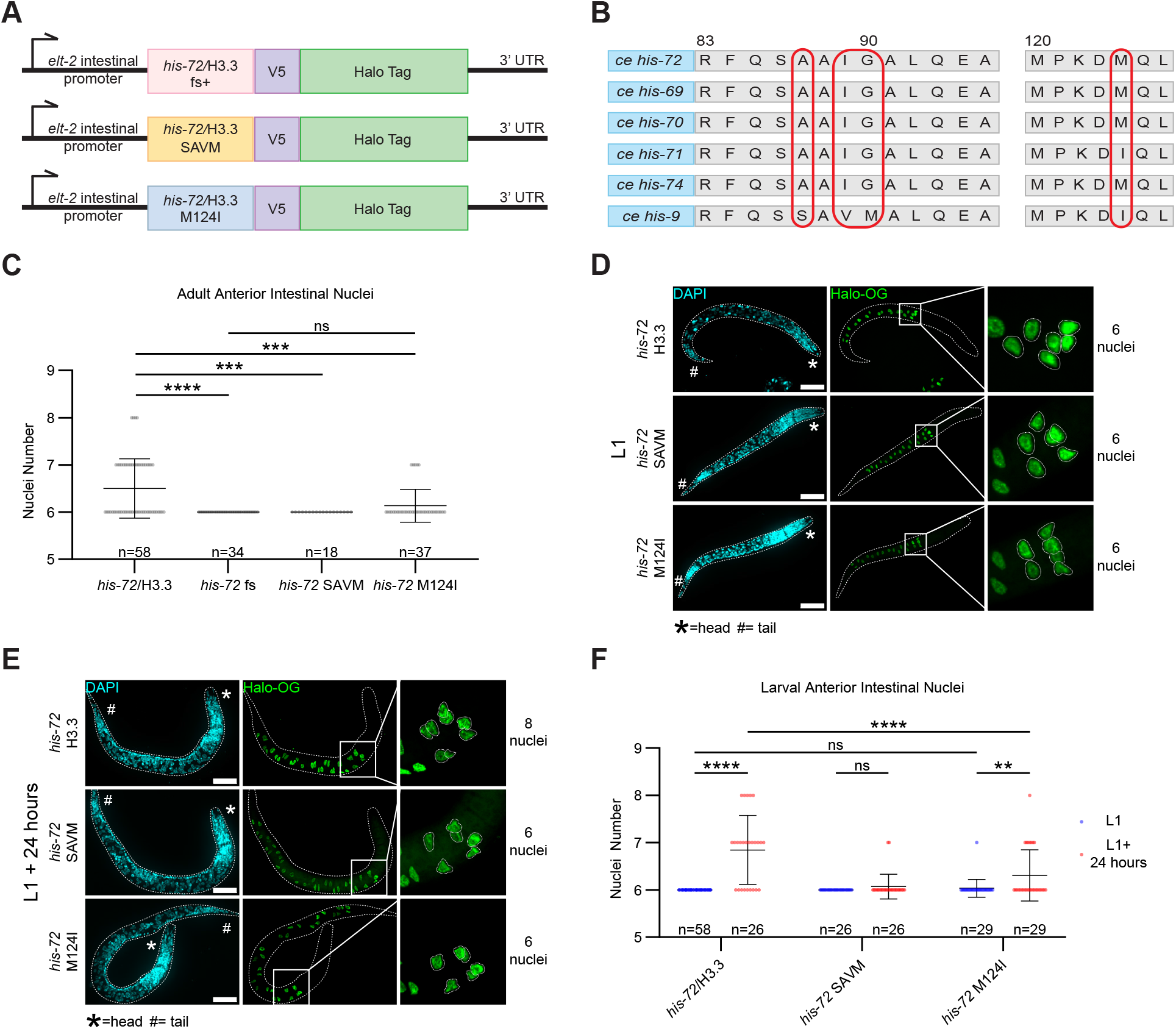
Mutations predicted to alter *his-72*/H3.3 incorporation ameliorate the intestinal extra nuclei phenotype. (**A**) Mutant *his-72*/H3.3 reporters generated. fs+, frame shift. (**B**) *C. elegans* histone H3 variants. *his-69, his-70, his-71, his-72, his-74* are histone H3.3 members. Red circles highlight amino acids critical for proper chromatin incorporation. (**C**) Anterior intestinal nuclei quantification of adult *his-72*/H3.3 reporter mutant worms. *his-72*/H3.3 adult data used in Fig 1D shown here. Significance determined by one-way ANOVA with Tukey’s multiple comparisons test; ***p<0.001; ****p<0.0001; ns, not significant. (**D-E**) Fluorescent images of his-72/H3.3, *his-72*/H3.3 SAVM, and *his-72*/H3.3 M124I at the first larval stage (D) or grown for 24 hours (E). Images shown as max intensity projection of confocal stack taken at 63X magnification under oil immersion. Anterior nuclei shown at either 5x (L1) or 3x (L1+24 hours) magnification from original image. *his-72*/H3.3 L1 and L1+24 hours images from Fig 2A, B shown here. Scale bar, 10 µm. (**F**) Quantification of *his-72*/H3.3 mutant worms collected at various developmental stages. L1 data shown in blue, while L1+24 hours data shown in red. *his-72*/H3.3 data from Fig 2D shown here. Significance determined by two-way ANOVA with Tukey’s multiple comparisons test; **p<0.01; ****p<0.0001; ns, not significant.

Histones must be deposited into chromatin to perform their molecular functions. Classically, histone H3.3 is recognized by the HIRA chaperone complex for chromatin deposition (7, 12). HIRA chaperone specificity for histone H3.3 is mediated by a conserved amino acid sequence motif that differs from the canonical histone H3.1 sequence (7) **(Fig 3B)**. To investigate HIRA chaperone involvement in intestinal extranuclear divisions, the *his-72*/H3.3 reporter sequence motif was mutated to the canonical histone H3.1 sequence **(Fig 3A, B;** see **Methods)**. The HIRA chaperone mutation (SAVM) significantly decreased the extranuclear phenotype in adults and in L1 worms developed for 24 hours **(Fig 3C-F; Fig S1E)**, implying deposition of *his-72*/H3.3 by the HIRA chromatin chaperone is critical for the extra nuclei phenotype. *C. elegans* have five H3.3 paralogs, *his-69, his-70, his-71, his-72*, and *his-74*, with an amino acid variation at position 124 of either isoleucine or methionine (19, 27) **(Fig 3B)**. H3 histone variants containing I124 have previously shown to reduce their incorporation into nucleosomes (28). CRISPR/Cas9 was used on *his-72*/H3.3 reporter worms to change the histone reporter sequence from methionine to isoleucine at amino acid 124 **(Fig 3A, B)**. The *his-72* M124I mutation significantly decreased the extranuclear phenotype in adults and in L1s developed for 24 hours **(Fig 3C-F; Fig S1E)**. Collectively, mutations of conserved histone H3.3 residues involved in chromatin incorporation were necessary for the extra nuclei phenotype in *his-72*/H3.3 reporter worms.

### Extra intestinal nuclei in H3.3 reporters contain lower chromosome copy number than undivided nuclei

Intestinal cells that underwent an extranuclear division contained nuclei that were smaller in size compared to undivided nuclei controls **(Fig S2A)**, suggesting potential defects in chromosome segregation or DNA replication during development. The anterior intestinal nuclei (cells 1-6) of developing larvae do not divide but undergo four rounds of endoreduplication at each larval molt, reaching 32C DNA content by adulthood (16) **(Fig 4A-I.)**. Inclusion of nuclear division in extra nuclei will cause 16C DNA content **(Fig 4A-II.)**. In contrast, central intestinal nuclei (cells 7-16) undergo mitosis without cytokinesis and four rounds of endoreduplication at each larval molt, generating nuclei with a 32C DNA content (16) **(Fig 4A-III.)**. Additionally, perturbations in nuclear mitotic division can lead to aneuploidy, or abnormal chromosome number, through defects in DNA replication, spindle microtubule fidelity, kinetochore binding, or chromatid segregation, which would produce nuclei with chromosome copy number variance in adults (29) **(Fig 4A-IV.)**.

**Fig 4.**
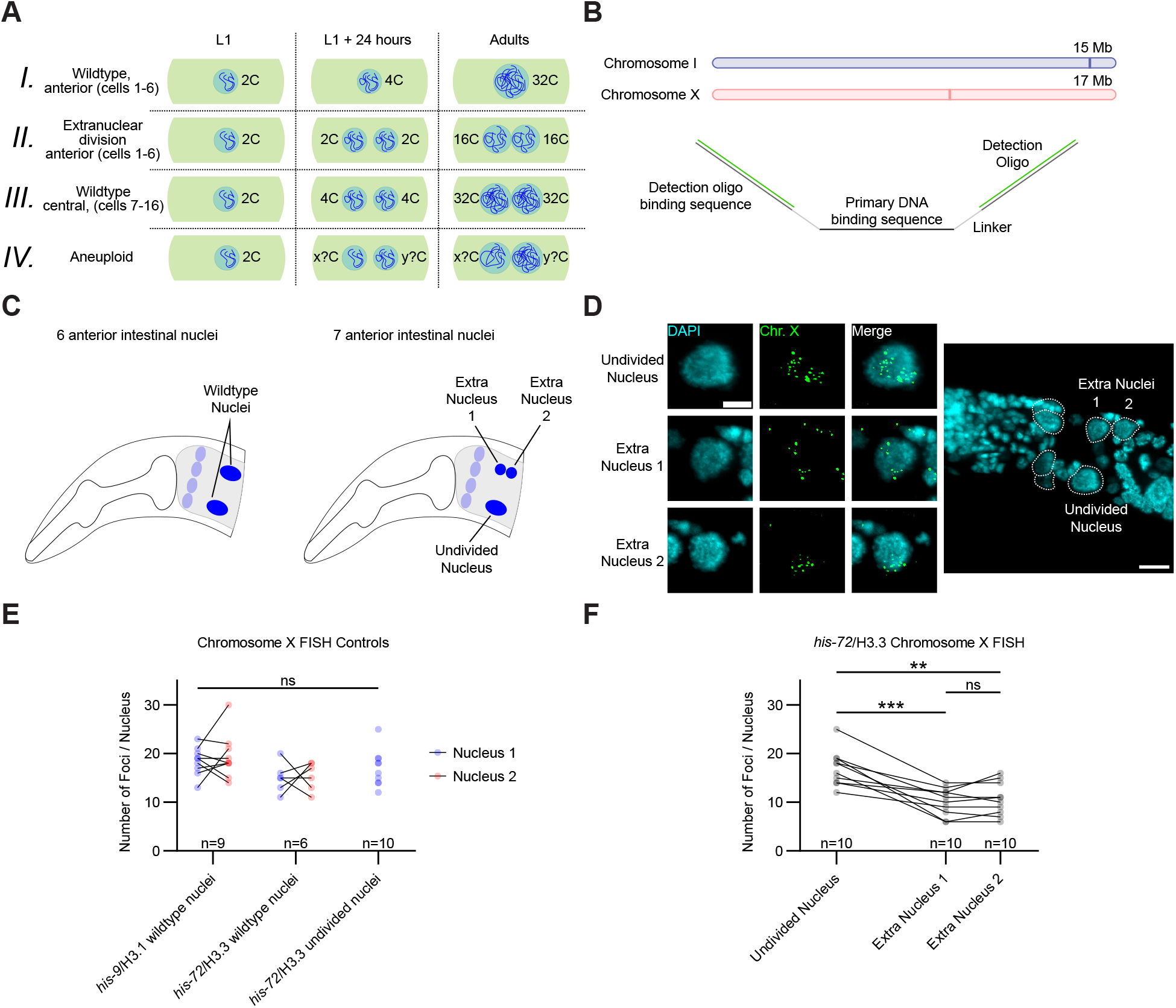
Extra nuclei divide normally and possess normal chromosome copy number. (**A**) Intestinal nuclei chromosome copy number throughout development. (I) Wild type anterior nuclei proceed through rounds of endoreduplication for 32C DNA content. (II) Including nuclear division in this process would cause DNA content to be divided between two nuclei. (III) In the central intestine, cells undergo mitosis without cytokinesis and rounds of endreduplication for two 32C nuclei. (IV) Aberrant chromosome segregation causes aneuploidy. (**B**) DNA Fluorescence In situ Hybridization (FISH) probe libraries target 50 kb regions on chromosome I and X. Probes consist of two detection oligo binding sites, two short linkers, and a 26-32 bp sequence complementary to the genome. Primary probes can hybridize to Alexa Fluor 488 detection oligos. (**C**) Schematic of anterior nuclei organization in wildtype (6 nuclei) and extra nuclei (7 nuclei) worms. (**D**) Representative images of a *his-72*/H3.3 reporter worm with abnormal 7 nuclei hybridized with chromosome X FISH oligos. Scale bar, 5 µm in nucleus image and 10 µm in worm image. (**E**) Comparison of counted foci in *his-9*/H3.1 and *his-72*/H3.3 wildtype and undivided nuclei hybridized with chromosome X FISH probes. Significance determined by two-way ANOVA with Tukey’s multiple comparisons test; ns, not significant. (**F**) Comparison of counted foci between undivided nuclei and the two divided extra nuclei in *his-72*/H3.3 worms possessing 7 anterior nuclei. Significance determined by repeated measures one-way ANOVA with Tukey’s multiple comparisons test; ***p<0.001, **p<0.01, ns, not significant.

To determine the chromosome state of extra nuclei, DNA Fluorescence In situ Hybridization (FISH) was performed to measure the copy number of two chromosomes in the anterior intestinal nuclei of adult worms. FISH oligonucleotide probes were designed to target chromosome I and X. They included a sequence for secondary fluorophore probe hybridization to facilitate visualization **(Fig 4B**, see **Methods)**. Synchronized adult worms were collected, hybridized with primary and secondary detection probes, and imaged (see **Methods)**. Fluorescent foci in *his-72*/H3.3 worms with 7 anterior nuclei were quantitated, as the chromosome copy number in undivided and extra-division nuclei could directly be compared in the same worm **(Fig 4C)**. *his-72*/H3.3 and *his-9*/H3.1 worms with the expected 6 anterior nuclei were also quantified as controls. As expected, there were no significant differences between chromosome copy number in *his-72*/H3.3 and *his-9*/H3.1 control nuclei **(Fig 4D, E; S2C)**. Chromosome copy number in *his-72*/H3.3 worms revealed significant differences between undivided and extra nuclei for both chromosomes I and X **(Fig 4D, F; Fig S2B, D)**. Comparison of copy number between extra nuclei revealed no significant difference and never exceeded 16 in count **(Fig 4F; S2D)**, implying defects in chromosome segregation during nuclear division causing aneuploidy are not likely. To support the FISH results, DNA content of divided and undivided anterior nuclei was measured in adult *his-72*/H3.3 reporters with seven anterior nuclei. Ventral cord neuronal nuclei served as an internal 2C control (30) (see **Methods)**. The mean DNA content of undivided nuclei controls was 32.57 +/− 5.06 compared to 14.87 +/− 3.340 and 17.01 +/− 3.360 for divided nuclei **(Fig S2E)**. Thus, the DNA content of extra nuclei was approximately half that of undivided nuclei. Collectively, these data support the model that anterior intestinal nuclei in *his-72*/H3.3 worms undergo division during the first 24 hours of larval development. These extra nuclei undergo rounds of endoreduplication for 16C DNA content in adult worms, half of what is found in undivided nuclei.

*his-72* is known to deposit into chromatin sites of active transcription and can maintain open chromatin conformations (19, 27). The *his-72*/H3.3 reporter examined here is atypically expressed through the *elt-2* promoter. This exogenous histone H3.3 expression may aberrantly express genes responsible for nuclear division. CDC-25.2 drives nuclear division in the *C. elegans* intestine by promoting cell cycle progression through activation of the CDK-1/CYB-1 cyclin-dependent kinase complex (31). *his-72*/H3.3 reporter function may disrupt the nuclear division axis through a similar mechanism. Similarly, overexpression of a human histone H3.3 variant, *H3f3a*, in lung cancer cells led to aberrant deposition and mis-expression of metastasis genes (32, 33). Investigating sites of *his-72*/H3.3 reporter chromatin deposition will connect gene expression changes to the extra nuclei phenotype observed.

Histone genes are often mutated in cancer (reviewed in (13, 34)). Dysregulation of histone H3.3 post translational modifications is a key driver of gliomas and other cancers (13, 35). The well-characterized H3.3(K27M) oncohistone mutation perturbs methylation of K27 by the PRC2 complex for increased acetylation (13, 35). Acetylation at this site increases chromatin accessibility, driving gene activation by recruitment by key transcriptional factors such as RNA polymerase II (13, 35). H3.3 is reported to accumulate in terminally differentiated cells of mice and is accompanied by changes in global histone methylation and genome organization (36, 37), illustrating the importance of H3.3 in mediating chromatin organization and gene expression. Future work will dissect chromatin post translational modifications and the role of chromatin architecture on gene regulation throughout *C. elegans* intestinal nuclei development.

Most histone mechanistic research has been performed either in cell culture or mitotic tissue. Although much has been learned, studies focusing on the role of histones in terminally differentiated tissues remains largely underexplored. H3.3 is known to accumulate in post mitotic neurons and is critical for establishing chromatin organization, the neuronal transcriptome, and ultimately cell identity (37). Histone H3.3 may be functioning similarly in the *C. elegans* intestine. Perturbations in H3.3 expression may alter chromatin organization, the transcriptome, and ultimately the function of the intestine tissue overall.

Endoreduplication is a hallmark of highly metabolic cell types and has been reported to occur in plant cells (38), human cardiomyocytes (39), hepatocytes (40), trophoblasts (41), keratinocytes (42), and megakaryocytes (43). In the *C. elegans* intestine, this process is essential for meeting the metabolic demands of the growing organism, as larger polyploid nuclei enable higher gene expression levels. Understanding how histones regulate nuclear dynamics in polyploid tissues could provide broader implications for developmental biology, tissue homeostasis, and even disease states where polyploidy is prevalent, such as cancer and liver regeneration.

## Materials and Methods

### Nematode Strains and Maintenance

Nematodes were cultured on NGM plates (25 mM KPO_4_ pH 6.0, 5 mM NaCl, 1 mM CaCl_2_, 1 mM MgSO_4_, 2.5 mg/ml tryptone, 5 µg/ml cholesterol, 1.7% agar) with HB101 bacteria as food source as described previously (44). All strains were grown from 15°C to 20°C. To synchronize populations at the first larval stage, worms were bleached and allowed to hatch overnight in M9 buffer using standard protocols (45). In temperature experiments, worms were exposed to specific temperatures as L4 larvae and allowed to propagate. F1 worms were collected, stained, and imaged as described below. The following strains were used in this study:

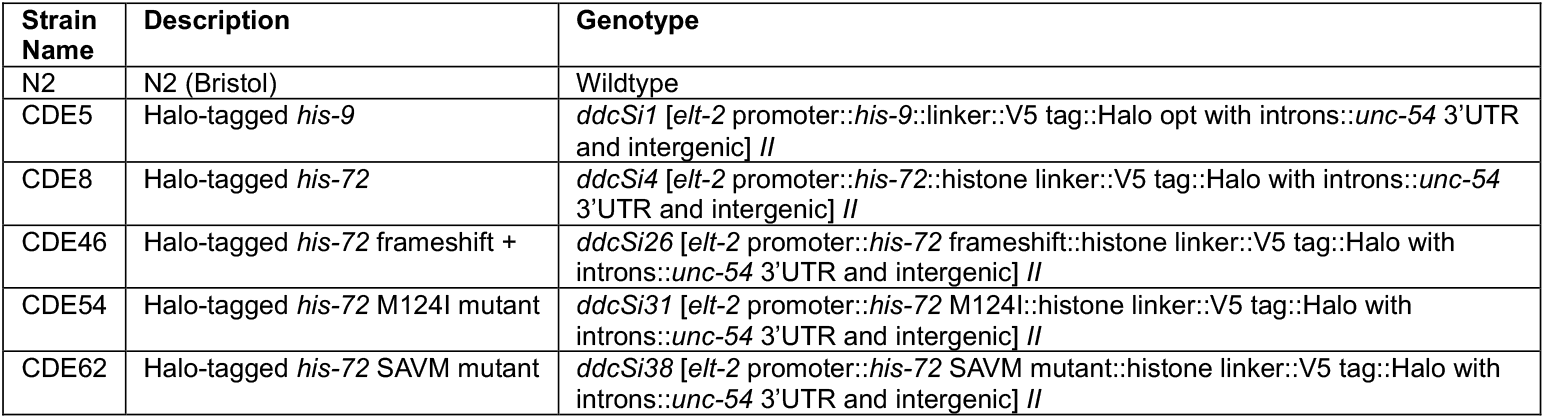

### CRISPR/Cas9 Gene Editin

A co-conversion CRISPR technique was implemented for targeted mutagenesis using a previously established protocol (46). Briefly, worms were injected with a target CRISPR/Cas9 RNA (crRNA) or plasmid expressing a Cas9-scaffold with tandem target sequence RNA (sgRNA) to a gene of interest, a target crRNA to *unc-58* or *dpy-10*, a scaffolding tracrRNA (IDT), recombinant Cas9 protein (IDT), and a *unc-58* or *dpy-10* repair DNA oligo that inserted a dominant mutation. F1s displaying the co-injection marker phenotype underwent additional screening by a combination of PCR without or with restriction enzyme digest to identify those with the repair of interest. F2s were PCR screened to identify homozygous alleles and the PCR product sequenced to confirm proper repair. The transgenic strain was crossed to the N2 strain for two generations to remove potential off target mutations. Endogenous *his-72* loci were sequenced after outcrossing to ensure wildtype sequences. All guide RNAs and oligos were obtained commercially (IDT). The primer sequences for crRNA and amplifying the repair template are listed below:

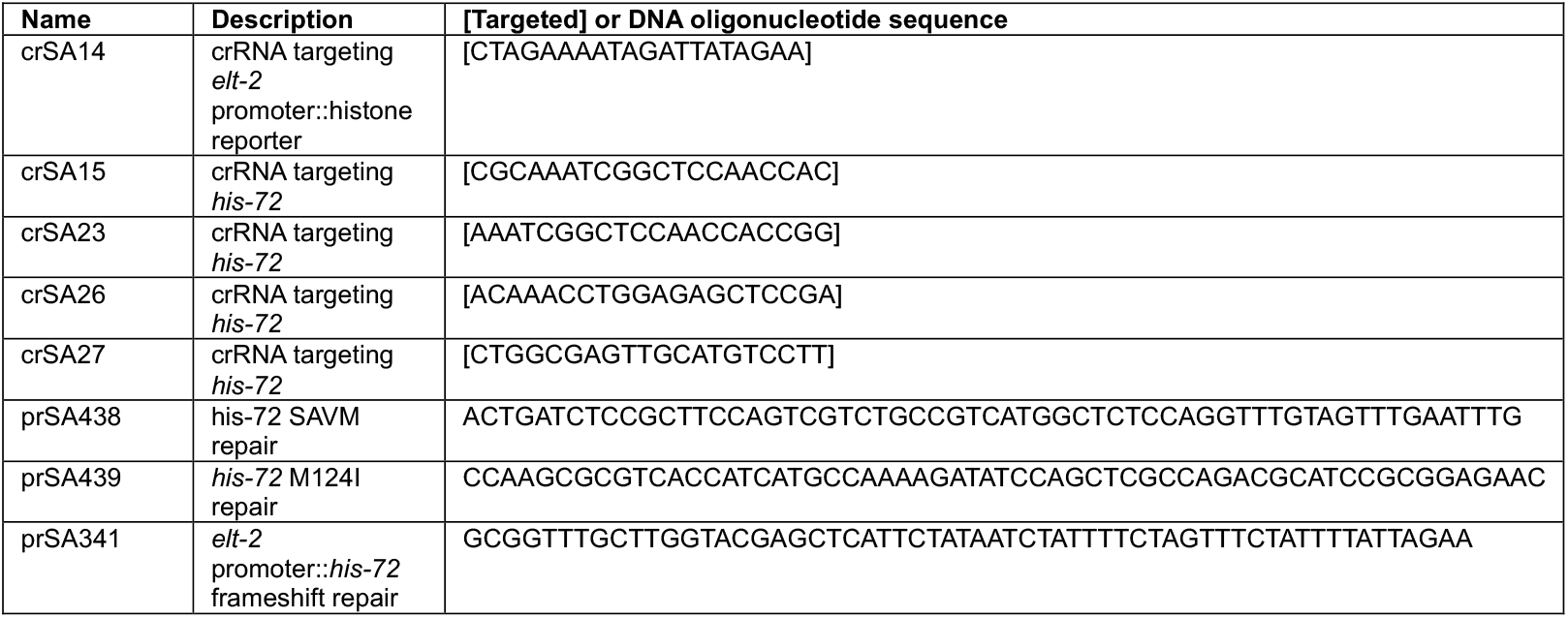

### Microscopy and image analysis

Worms were collected from agar plates into tubes with 1x Phosphate Buffer Saline (PBS; 137 mM NaCl, 2.7 mM KCl, 10 mM Na_2_HPO_4_, 1.8 mM KH_2_PO_4_) + 0.01% Tween-20 (PBS-T), washed three times to remove excess bacteria, fixed for 10 minutes with 3% paraformaldehyde (PFA) (EMS, 50-980-487), and stained with DAPI and Halo-Oregon Green (Promega) for 1 hour. Stained worms were mounted onto a slide with Vectashield (H-1900-10, Vector Laboratories Inc; Newark, CA) and imaged on a Zeiss AxioObserverZ1 modified by 3i (www.intelligent-imaging.com) for confocal microscopy. All images were analyzed by ImageJ/Fiji (47). Within any set of comparable images, the image capture and scaling conditions were kept identical for reproducibility. The cyan/blue channel included a 405-emission filter, and a green channel included a 488**-**emission filter. For each reporter, the anterior set or all intestinal nuclei were analyzed. A minimum of two biological replicates were performed. DNA content measurements of intestinal nuclei was performed as previously described (30). Briefly, DNA content values were estimated by determining the DAPI-based densitometric quantification and normalizing to 2C ventral cord nuclei.

### Immunoblot Analysis

Synchronized populations of *C. elegans* adult hermaphrodites were harvested, washed three times in PBS-T, and stored in −20°C with 5x SDS buffer (250 mM Tris-HCl pH 6.8, 25 mM EDTA pH 8, 25% glycerol, 5% SDS). The samples were then thawed, boiled for 10 minutes at 90°C, centrifuged for 1 minute at 10,000 rpm and loaded onto SDS-PAGE gels. Proteins were transferred to a PVDF membrane (BioRad, #1620264) using the BioRad TransBlot Turbo System (BioRad, #1704150). Membranes were blocked for 1 hour at room temperature with PBS-T buffer containing 5% non-fat milk and incubated for 2 hours at room temperature with HRP-conjugated antibodies. Membranes were then washed 3 times with PBS-T. The following antibodies or protein-HRP conjugates were used for this study: V5-Tag (E9H8O) Mouse mAb #80076 (1:1000) (Cell Signaling), and GAPDH (D4C6R) Mouse mAb #97166 (1:5000) (Cell Signaling).

### DNA FISH

#### Probe Design

Primary probe sequences targeting 50 kb regions on chromosome I and X were selected from the Wu Lab’s Oligopaint FISH oligonucleotide repository (https://oligopaints.hms.harvard.edu/) using the *C. elegans* genome (*ce11*) and were identified using the ‘coverage’ mining setting of the Oligominer program (48). A short linker and detection oligo binding sequence were appended to the 5’ and 3’ end of primary probe sequences as designed by others (49). Probe sets were generated (IDT) targeting two chromosomes, one autosomal (I) and one sex (X) to ensure consistency and reproducibility of results. (**Supplemental File 1**). The detection probe sequence were also designed by others (49) and modified on the 5’ end with an Alexa Fluor 488 fluorophore (IDT) (**Supplemental File 1**).

#### FISH Protocol

The DNA fluorescence in situ hybridization (FISH) protocol was performed as described by Fields et al. (49) in adult worms with the following modifications. Worms were maintained on NGM plates seeded with *E. coli* HB101 until the bacterial lawn was almost exhausted. They were then washed off plates with M9 medium (42.3 mM Na_2_HPO_4_, 22 mM KH_2_PO_4_, 85.6 mM NaCl, 1 mM MgSO_4_) into 15 mL conical tubes. Samples were bleached using standard protocols and allowed to hatch overnight in M9 buffer to synchronize populations at the first larval stage (45). Synchronized L1s were moved to NGM plates and allowed to develop for 72 hours. Young adult worms were collected and washed with M9 to remove all excess bacteria. Worms were fixed in 3% PFA for 10 minutes followed by storage at −20°C in 100% ice cold methanol until use. 100 pmol of chromosome I and X primary probe pools were hybridized overnight at 37°C. Unbound probe was washed away with 2x SSC-T (Fisher Scientific) and samples incubated with 100 pmol detection probe for 3 hours at room temperature. For DNA FISH image analysis, the 488 channel was set to the same fluorescent levels for all images. A rectangle was drawn around individual nuclei and the number of z-plane stacks was determined. This was used to crop the image to only include individual intestinal nuclei. The 488 channel was duplicated twice, and the Gaussian filter set to 1.0 and 2.0, respectively. Image calculator was used to subtract the 1.0 Gaussian filtered image from the 2.0 image to improve the foci signal-to-noise. Foci were counted using the 3D objects counter in ImageJ FIJI. The threshold and image size were maintained between all images. Images with less than 10 foci in nuclei were excluded. Area of nuclei was determined by using fluorescence thresholding to mark the intestinal nuclei and using the area function. FISH experiments were performed at least three times.

## Acknowledgements

The authors thank members of the Aoki lab for thoughtful discussions. B.B. was supported by the Indiana Medical Scientist Training Program (MSTP) summer research program. S.T.A. and this work was funded by the Indiana University Precision Health Initiative, Showalter Trust, and NIH/NIGMS (R35 GM142691).

## Supplementary Figure captions

**Fig S1.**
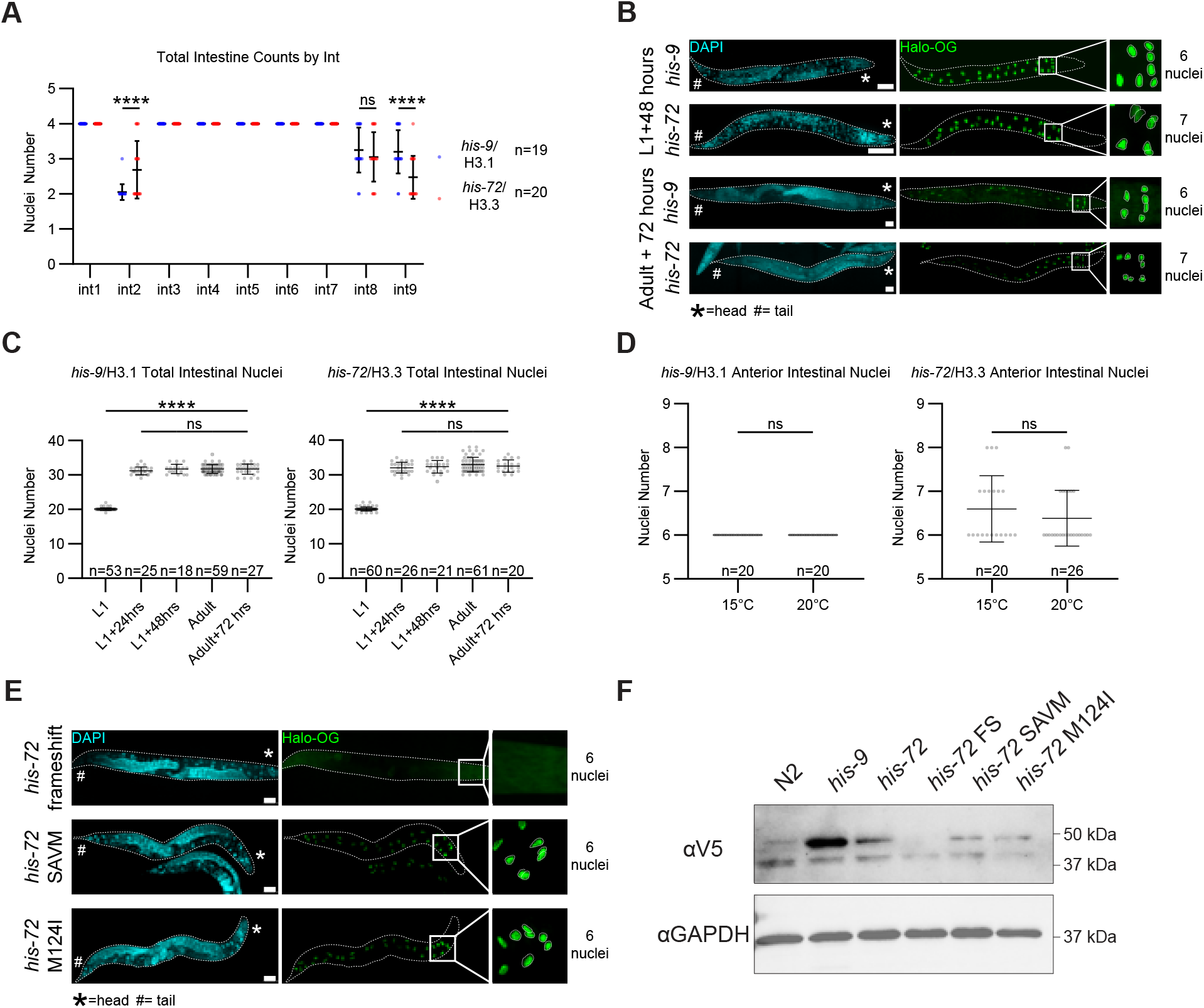
Additional results of the intestinal extra nuclei phenotype. (**A**) Nuclei counts in each int of the intestine. Significance determined by two-way ANOVA with Tukey’s multiple comparisons test; ****p<0.0001; ns, not significant. (**B**) Fluorescent images of *his-9*/H3.1 and *his-72*/H3.3 reporters at various developmental stages. Images shown as a combined montage max intensity projection of confocal stack taken at 40X magnification under oil immersion. (**C**) Total intestinal nuclei number from *his-9*/H3.1 and *his-72/*H3.3 worms are various developmental stages. Significance determined by one-way ANOVA with Tukey’s multiple comparisons test; ****p<0.0001; ns, not significant. (**D**) Anterior nuclei number in *his-9*/H3.1 and *his-72*/H3.3 adult worms cultured at 15°C or 20°C. Significance determined by unpaired students t-test; ns, not significant. (**E**) Fluorescent images of *his-72* frameshift, *his-72* SAVM, and *his-72* M124I adult worms. Images shown as a combined montage max intensity projection of confocal stack taken at 40X magnification under oil immersion. (**F**) Immunoblot of *his-72/*H3.3 mutants from 100 synchronized adult worms. Additional band of lower molecular weight was consistently present across all experimental conditions and is likely background. Reporter proteins on the same blot were detected by V5 and GAPDH loading control antibodies.

**Fig S2.**
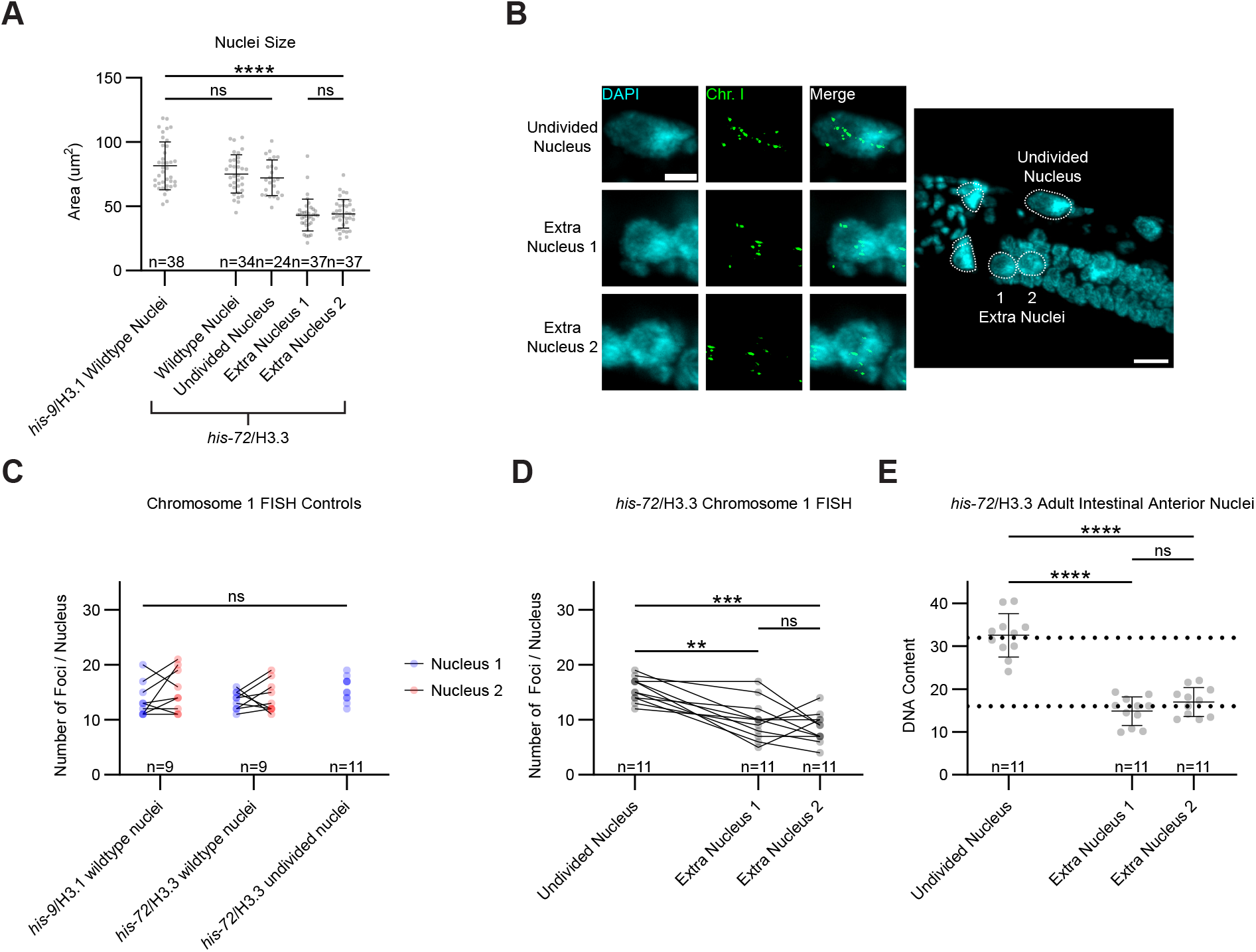
Additional chromosome quantification results in the intestine. (**A**) Area of anterior nuclei in *his-72*/H3.3 and *his-9*/H3.1 reporter worms. Significance determined by one-way ANOVA with Tukey’s multiple comparisons test; ****p<0.0001; ns, not significant. (**B**) Representative images of *his-72*/H3.3 worms with abnormal 7 nuclei hybridized with chromosome I FISH probes. Scale bar, 5 µm in nucleus image and 10 µm in worm image. (**C**) Comparison of counted foci in *his-9*/H3.1 and *his-72*/H3.3 wildtype and undivided nuclei hybridized with chromosome I FISH probes. Significance determined by two-way ANOVA with Tukey’s multiple comparisons test; ns, not significant. (**D**) Comparison of counted foci between undivided nuclei and the two divided extra nuclei in *his-72*/H3.3 worms possessing 7 anterior nuclei. Significance determined by repeated measures one-way ANOVA with Tukey’s multiple comparisons test; ***p<0.001, **p<0.01, ns, not significant. (**E**) DNA content of undivided and extra nuclei in *his-72*/H3.3 worms with abnormal seven anterior nuclei. Dotted lines mark 32C and 16C. Significance determined by repeated measures one-way ANOVA with Tukey’s multiple comparisons test; ****p<0.0001, ns, not significant.

## References

1. Olins, A. L., and Olins, D. E. (1974) Spheroid Chromatin Units (v Bodies) Science 183, 330–332

2. Kornberg, R. D. (1974) Chromatin Structure: A Repeating Unit of Histones and DNA Science 184, 868–871

3. Kornberg, R. D., and Thomas, J. O. (1974) Chromatin Structure: Oligomers of the Histones Science 184, 865–868

4. Luger, K., Mäder, A. W., Richmond, R. K., Sargent, D. F., and Richmond, T. J. (1997) Crystal structure of the nucleosome core particle at 2.8 Å resolution Nature 389, 251–260

5. Martire, S., and Banaszynski, L. A. (2020) The roles of histone variants in fine-tuning chromatin organization and function Nature Reviews Molecular Cell Biology 21, 522–541

6. Smith, S., and Stillman, B. (1989) Purification and characterization of CAF-I, a human cell factor required for chromatin assembly during DNA replication in vitro Cell 58, 15–25

7. Tagami, H., Ray-Gallet, D., Almouzni, G., and Nakatani, Y. (2004) Histone H3.1 and H3.3 Complexes Mediate Nucleosome Assembly Pathways Dependent or Independent of DNA Synthesis Cell 116, 51–61

8. Nakano, S., Stillman, B., and Horvitz, H. R. (2011) Replication-Coupled Chromatin Assembly Generates a Neuronal Bilateral Asymmetry in C. elegans Cell 147, 1525–1536

9. Harada, A., Maehara, K., Sato, Y., Konno, D., Tachibana, T., Kimura, H. et al. (2015) Incorporation of histone H3.1 suppresses the lineage potential of skeletal muscle Nucleic Acids Res 43, 775–786

10. Stroud, H., Otero, S., Desvoyes, B., Ramírez-Parra, E., Jacobsen, S. E., and Gutierrez, C. (2012) Genome-wide analysis of histone H3.1 and H3.3 variants in Arabidopsis thaliana Proceedings of the National Academy of Sciences 109, 5370–5375

11. Goldberg, A. D., Banaszynski, L. A., Noh, K.-M., Lewis, P. W., Elsaesser, S. J., Stadler, S. et al. (2010) Distinct Factors Control Histone Variant H3.3 Localization at Specific Genomic Regions Cell 140, 678–691

12. Burkhart, K. B., Sando, S. R., Corrionero, A., and Horvitz, H. R. (2020) H3.3 Nucleosome Assembly Mutants Display a Late-Onset Maternal Effect Current Biology 30, 2343–2352.e2343

13. Wong, L. H., and Tremethick, D. J. (2024) Multifunctional histone variants in genome function Nature Reviews Genetics 26, 82–104

14. Dimov, I., and Maduro, M. F. (2019) The C. elegans intestine: organogenesis, digestion, and physiology Cell and Tissue Research 377, 383–396

15. Sulston, J. E., and Horvitz, H. R. (1977) Post-embryonic cell lineages of the nematode, Caenorhabditis elegans Developmental Biology 56, 110–156

16. Hedgecock, E. M., and White, J. G. (1985) Polyploid tissues in the nematode Caenorhabditis elegans Developmental Biology 107, 128–133

17. Borchers, C., Osburn, K., Roh, H. C., and Aoki, S. T. (2025) In vivo pulse-chase in C. elegans reveals intestinal histone turnover changes upon starvation bioRxiv 10.1101/2025.02.13.638128

18. Postberg, J., Forcob, S., Chang, W.-J., and Lipps, H. J. (2010) The evolutionary history of histone H3 suggests a deep eukaryotic root of chromatin modifying mechanisms BMC Evolutionary Biology 10,

19. Ooi, S. L., Priess, J. R., and Henikoff, S. (2006) Histone H3.3 Variant Dynamics in the Germline of Caenorhabditis elegans PLoS Genetics 2,

20. Fukushige, T., Hawkins, M. G., and McGhee, J. D. (1998) The GATA-factor elt-2 is essential for formation of the Caenorhabditis elegans intestine Developmental Biology 198, 286–302

21. McGhee, J. D., Fukushige, T., Krause, M. W., Minnema, S. E., Goszczynski, B., Gaudet, J. et al. (2009) ELT-2 is the predominant transcription factor controlling differentiation and function of the C. elegans intestine, from embryo to adult Developmental Biology 327, 551–565

22. Los, G. V., Encell, L. P., McDougall, M. G., Hartzell, D. D., Karassina, N., Zimprich, C. et al. (2008) HaloTag: A Novel Protein Labeling Technology for Cell Imaging and Protein Analysis ACS Chemical Biology 3, 373–382

23. McGee, M. D., Weber, D., Day, N., Vitelli, C., Crippen, D., Herndon, L. A. et al. (2011) Loss of intestinal nuclei and intestinal integrity in aging C. elegans Aging Cell 10, 699–710

24. Chen, L., Wang, Y., Zhou, X., Wang, T., Zhan, H., Wu, F. et al. (2023) Investigation into the communication between unheated and heat-stressed Caenorhabditis elegans via volatile stress signals Scientific Reports 13,

25. Jovic, K., Sterken, M. G., Grilli, J., Bevers, R. P. J., Rodriguez, M., Riksen, J. A. G. et al. (2017) Temporal dynamics of gene expression in heat-stressed Caenorhabditis elegans PLoS One 12, e0189445

26. Kyriakou, E., Taouktsi, E., and Syntichaki, P. (2022) The Thermal Stress Coping Network of the Nematode Caenorhabditis elegans Int J Mol Sci 23,

27. Delaney, K., Mailler, J., Wenda, J. M., Gabus, C., and Steiner, F. A. (2018) Differential Expression of Histone H3.3 Genes and Their Role in Modulating Temperature Stress Response in Caenorhabditis elegans Genetics 209, 551–565

28. Kujirai, T., Horikoshi, N., Xie, Y., Taguchi, H., and Kurumizaka, H. (2017) Identification of the amino acid residues responsible for stable nucleosome formation by histone H3.Y Nucleus 8, 239–248

29. Potapova, T., and Gorbsky, G. J. (2017) The Consequences of Chromosome Segregation Errors in Mitosis and Meiosis Biology (Basel) 6,

30. Liu, M., Liu, P., Zhang, L., Cai, Q., Gao, G., Zhang, W. et al. (2011) mir-35 is involved in intestine cell G1/S transition and germ cell proliferation in C. elegans Cell Research 21, 1605–1618

31. Lee, Y.-U., Son, M., Kim, J., Shim, Y.-H., and Kawasaki, I. (2016) CDC-25.2, aC. elegansortholog ofcdc25, is essential for the progression of intestinal divisions Cell Cycle 15, 654–666

32. Park, S.-M., Choi, E.-Y., Bae, M., Kim, S., Park, J. B., Yoo, H. et al. (2016) Histone variant H3F3A promotes lung cancer cell migration through intronic regulation Nature Communications 7,

33. Klein, R. H., and Knoepfler, P. S. (2023) Knockout tales: the versatile roles of histone H3.3 in development and disease Epigenetics & Chromatin 16,

34. Lowe, B. R., Maxham, L. A., Hamey, J. J., Wilkins, M. R., and Partridge, J. F. (2019) Histone H3 Mutations: An Updated View of Their Role in Chromatin Deregulation and Cancer Cancers 11,

35. Saratsis, A. M., Knowles, T., Petrovic, A., and Nazarian, J. (2024) H3K27M mutant glioma: Disease definition and biological underpinnings Neuro-Oncology 26, S92–S100

36. Tvardovskiy, A., Schwämmle, V., Kempf, S. J., Rogowska-Wrzesinska, A., and Jensen, O. N. (2017) Accumulation of histone variant H3.3 with age is associated with profound changes in the histone methylation landscape Nucleic Acids Research 45, 9272–9289

37. Funk, O. H., Qalieh, Y., Doyle, D. Z., Lam, M. M., and Kwan, K. Y. (2022) Postmitotic accumulation of histone variant H3.3 in new cortical neurons establishes neuronal chromatin, transcriptome, and identity Proceedings of the National Academy of Sciences 119,

38. Tourdot, E., Mauxion, J.-P., Gonzalez, N., Chevalier, C., and Osorio, S. (2023) Endoreduplication in plant organogenesis: a means to boost fruit growth Journal of Experimental Botany 74, 6269–6284

39. Mollova, M., Bersell, K., Walsh, S., Savla, J., Das, L. T., Park, S.-Y. et al. (2013) Cardiomyocyte proliferation contributes to heart growth in young humans Proceedings of the National Academy of Sciences 110, 1446–1451

40. Sanjeev, G. (2000) Hepatic polyploidy and liver growth control Seminars in Cancer Biology 10, 161–171

41. Ouseph, Madhu M., Li, J., Chen, H.-Z., Pécot, T., Wenzel, P., Thompson John C. et al. (2012) Atypical E2F Repressors and Activators Coordinate Placental Development Developmental Cell 22, 849–862

42. Gandarillas, A. (2014) The mysterious human epidermal cell cycle, or an oncogene-induced differentiation checkpoint Cell Cycle 11, 4507–4516

43. Ravid, K., Lu, J., Zimmet, J. M., and Jones, M. R. (2001) Roads to polyploidy: The megakaryocyte example Journal of Cellular Physiology 190, 7–20

44. Brenner, S. (1974) The Genetics of Caenorhabditis Elegans Genetics 77, 71–94

45. Stiernagle, T. (2006) Maintenance of C. elegans WormBook 10.1895/wormbook.1.101.1

46. Paix, A., Folkmann, A., Rasoloson, D., and Seydoux, G. (2015) High Efficiency, Homology-Directed Genome Editing in Caenorhabditis elegans Using CRISPR-Cas9 Ribonucleoprotein Complexes Genetics 201, 47–54

47. Schindelin, J., Arganda-Carreras, I., Frise, E., Kaynig, V., Longair, M., Pietzsch, T. et al. (2012) Fiji: an open-source platform for biological-image analysis Nature Methods 9, 676–682

48. Beliveau, B. J., Kishi, J. Y., Nir, G., Sasaki, H. M., Saka, S. K., Nguyen, S. C. et al. (2018) OligoMiner provides a rapid, flexible environment for the design of genome-scale oligonucleotide in situ hybridization probes Proc Natl Acad Sci U S A 115, E2183–E2192

49. Fields, B. D., Nguyen, S. C., Nir, G., and Kennedy, S. (2019) A multiplexed DNA FISH strategy for assessing genome architecture in Caenorhabditis elegans eLife 8,

